## Supplementary figures and tables for "Mutationathon: towards standardization in estimates of pedigree-based germline mutation rates"

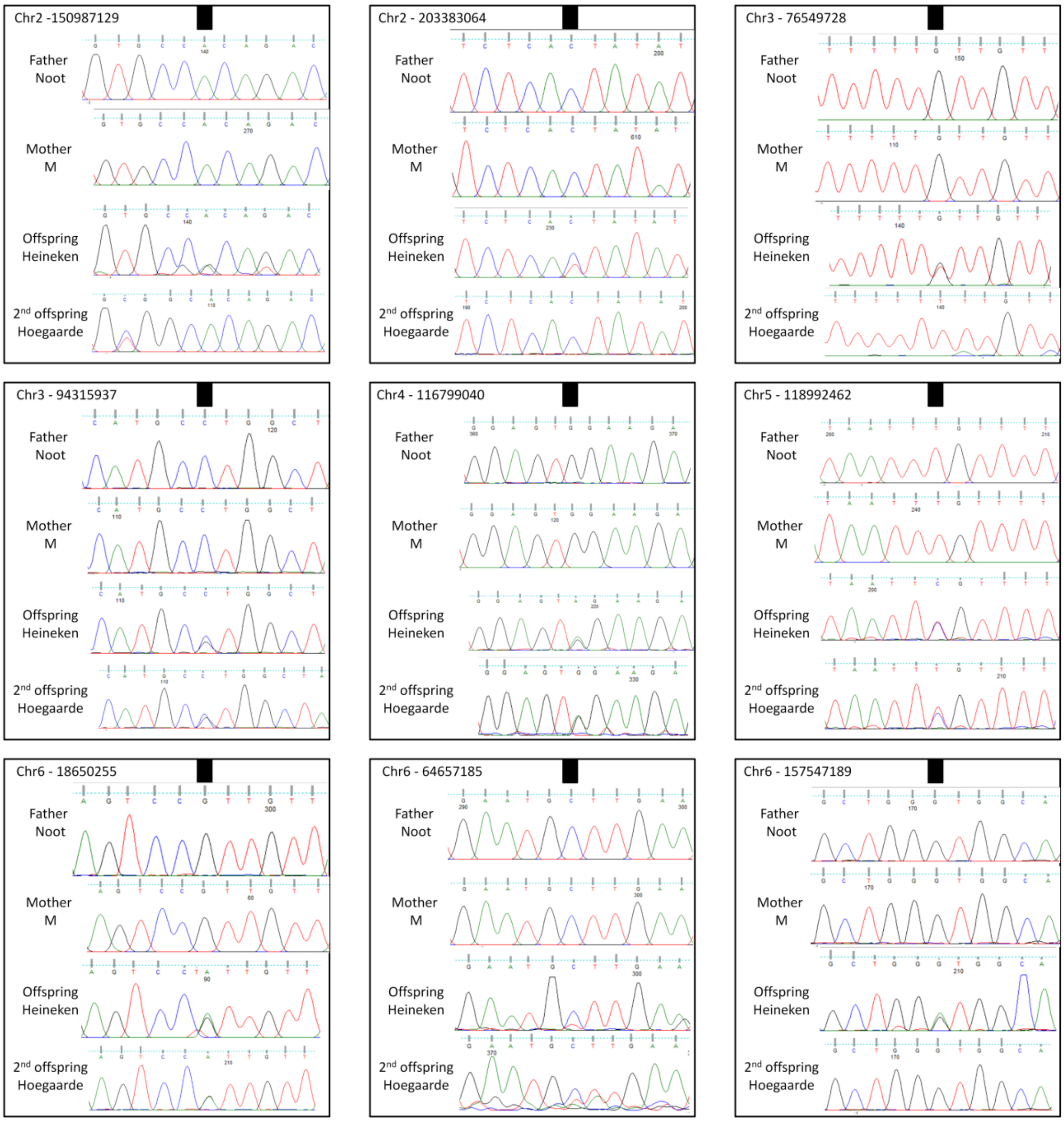

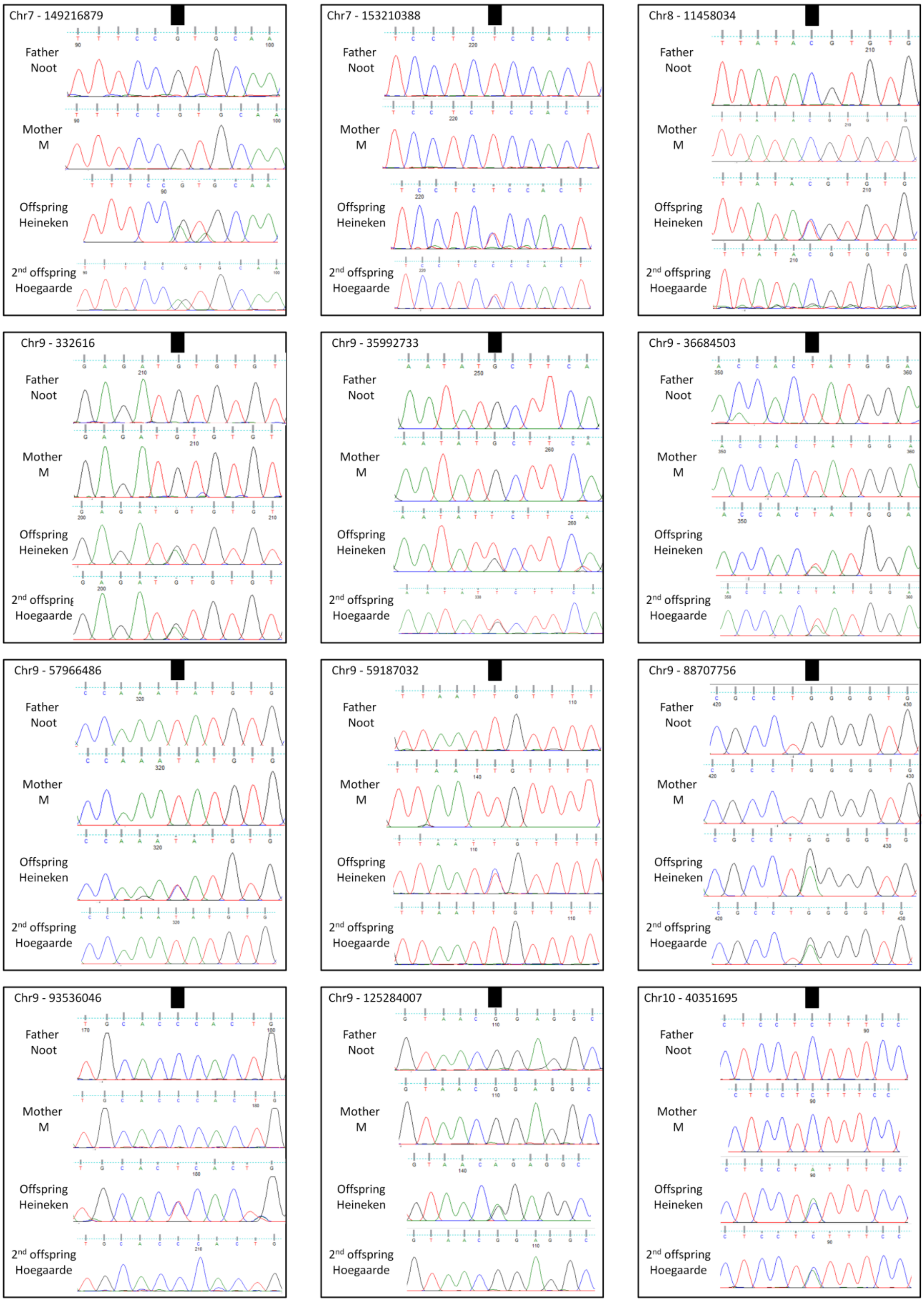
Chr7 - 149216879
Chr7 - 153210388
Chr8 - 11458034
Chr9 - 332616
Chr9 - 35992733
Chr9 - 36684503
Chr9 - 57966486
Chr9 - 59187032
Chr9 - 88707756
Chr9 - 93536046
Chr9 - 125284007
Chr10 - 40351695
Father Noot
Mother M
Offspring Heineken
2nd offspring Hoegaarde

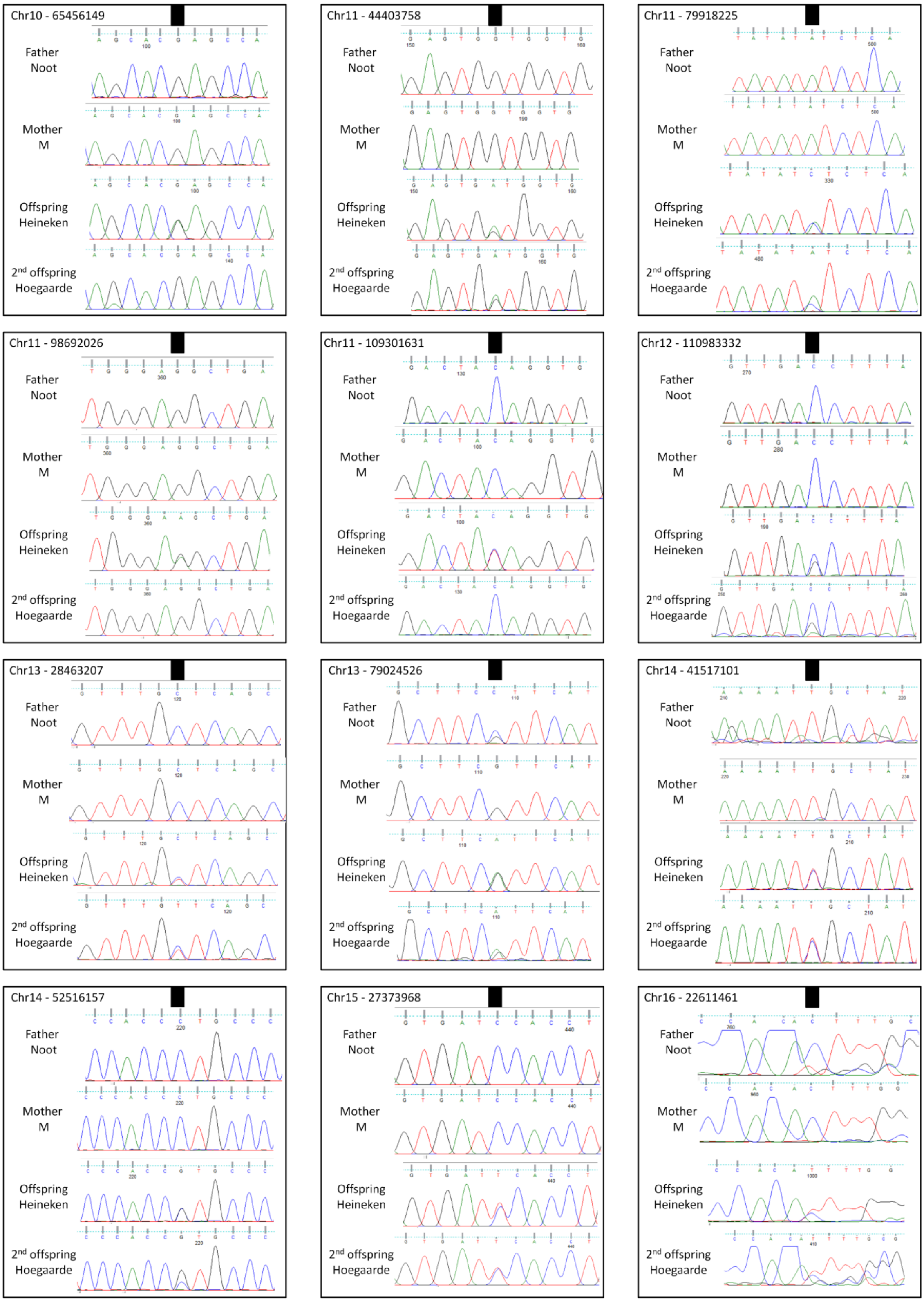

Chr10 - 65456149
Father Noot
Mother M
Offspring Heineken
2nd offspring Hoegaarde
Chr11 - 44403758
Father Noot
Mother M
Offspring Heineken
2nd offspring Hoegaarde
Chr11 - 79918225
Father Noot
Mother M
Offspring Heineken
2nd offspring Hoegaarde
Chr11 - 98692026
Father Noot
Mother M
Offspring Heineken
2nd offspring Hoegaarde
Chr11 - 109301631
Father Noot
Mother M
Offspring Heineken
2nd offspring Hoegaarde
Chr12 - 110983332
Father Noot
Mother M
Offspring Heineken
2nd offspring Hoegaarde
Chr13 - 28463207
Father Noot
Mother M
Offspring Heineken
2nd offspring Hoegaarde
Chr13 - 79024526
Father Noot
Mother M
Offspring Heineken
2nd offspring Hoegaarde
Chr14 - 41517101
Father Noot
Mother M
Offspring Heineken
2nd offspring Hoegaarde
Chr14 - 52516157
Father Noot
Mother M
Offspring Heineken
2nd offspring Hoegaarde
Chr15 - 27373968
Father Noot
Mother M
Offspring Heineken
2nd offspring Hoegaarde
Chr16 - 22611461
Father Noot
Mother M
Offspring Heineken
2nd offspring Hoegaarde

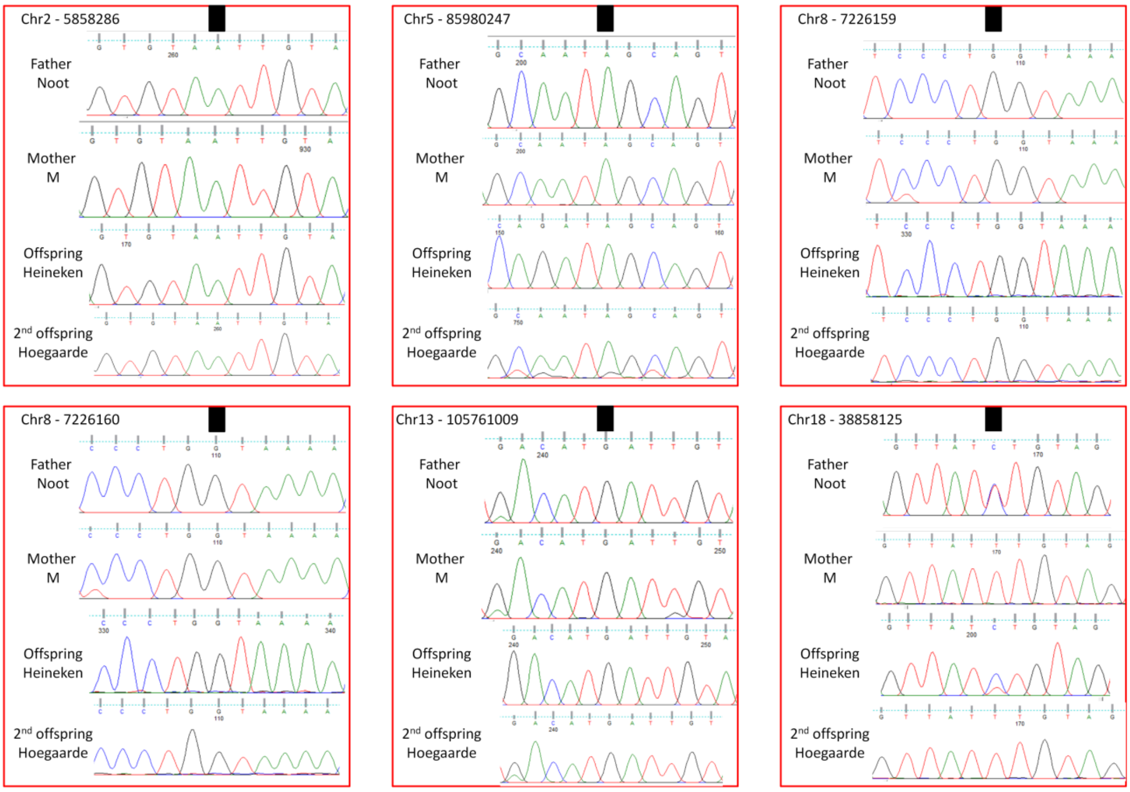

**Supplementary Figure 1 – Sanger sequencing chromatograms of the 39 DNM candidate sites that were successfully amplified for the four individuals, i.e. father (Noot), mother (M), offspring (Heineken), and second-generation offspring (Hoegaarde).** For each alignment, the candidate germline mutation position is located under the black square. The last six chromatograms (surrounded by red boxes) are the candidates that were detected as false-positive candidates.

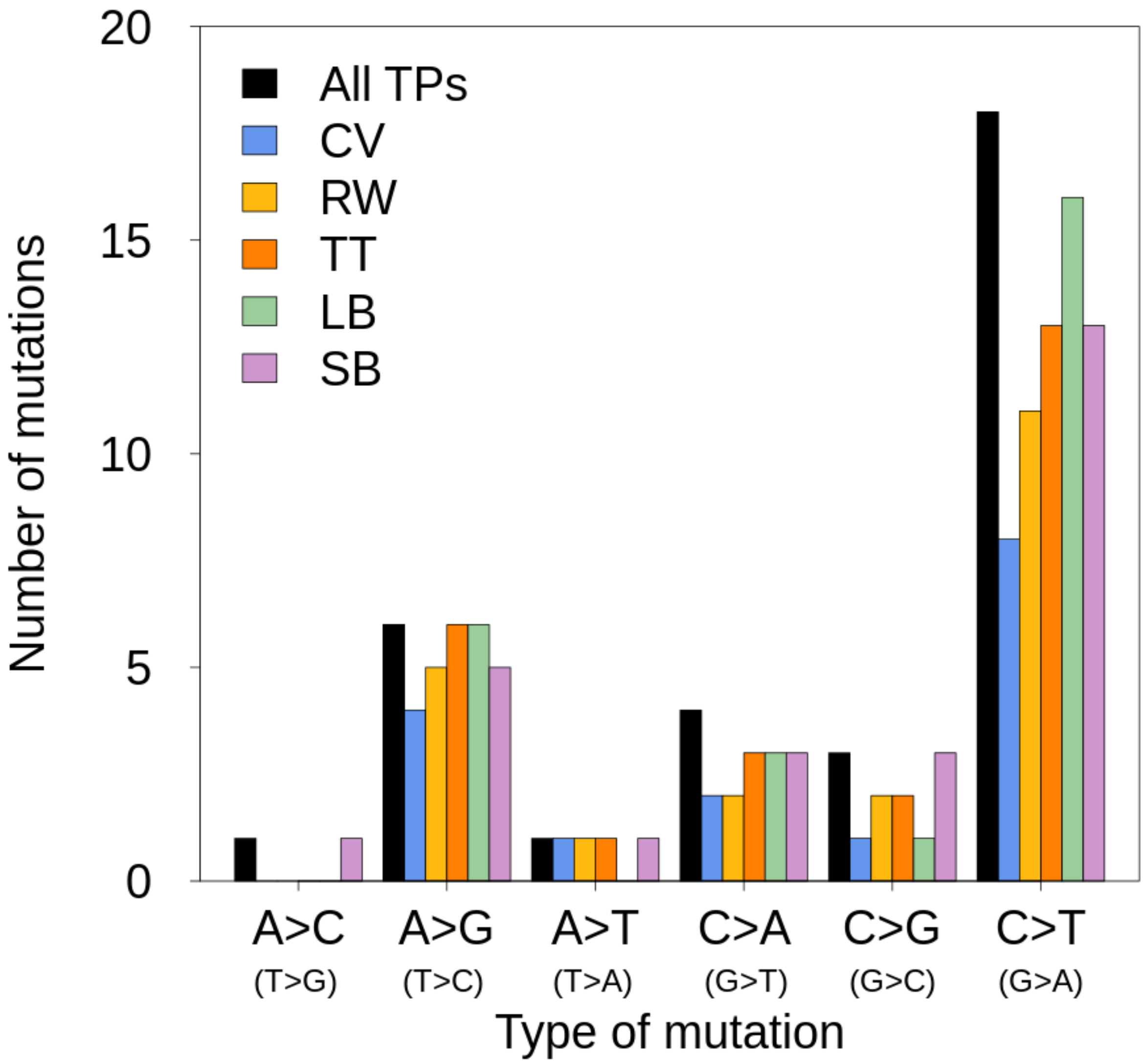

**Supplementary Figure 2 – Mutation spectrum of the trio of rhesus macaques.** "All TPs" corresponds to all true positive DNMs validated by the PCR experiment. The different colors correspond to the true positive DNMs found by each pipeline (LB: Lucie Bergeron, SB: Søren Besenbacher, CV: Cyril Versoza, TT: Tychele Turner, and RW: Richard Wang).

**Supplementary Table 1 – Study design and methodology of the studies on pedigree-based germline mutation rate estimation**. Information on the species studied, the number of trios, type of library preparation, the sequencing machine, sequencing depth, software used for mapping, variant calling, and filters used to detect the DNMs and calculate mutation rates are reported when available. Only recent studies from 2016 onwards are included.

| **Study** | **Species** | **Trios** | **Library** | **Sequencing** | **Depth** | **Mapping** | **Remove duplicates** | **BQSR** | **Variant calling** | **DNMs detection** | **Method FDR** | **FDR** | **Method FNR** | **FNR** | **Method *CG*** | ***CG*** | **$\mu$ per generation** |
|---|---|---|---|---|---|---|---|---|---|---|---|---|---|---|---|---|---|
| **Rahbari et al. 2016** | Human | 13 | | | 24.7X | | | | | DeNovoGear | Illumina resequencing and manual curation | | | | Remove sites not passing the DP filter, in low complex region, not validated by primers, or filtered by DeNovoGear priors filters | 83.1 % | Poisson distribution |
| **Smeds et al. 2016** | Collared flycatcher | 7 | | Illumina HiSeq | 40X | BWA 0.7.5a | Picard MarkDuplicates | Yes | GATK 3.3.0 - HaplotypeCaller and GenotypeGVCFs | Customized filtering | Manual curation | 35 % | | 0 | | 80 % | |

| | | | | | | | | | | | | | | | | | |
|---|---|---|---|---|---|---|---|---|---|---|---|---|---|---|---|---|---|
| **Wong et al. 2016** | Human | 719 | | Complete Genomics Inc. and 61 also with Illumina | 60X | Complete Genomics' Assembly (CGA) Pipeline versions 2.0.0–2.0.4 | | | CGA or Strelka and GATK | Customized filter and PhaseByTransmission | Overlap between different pipeline and different sequencing | 13 % | Overlap between different pipeline and different sequencing | 25 % | Remove from the commnly callable region the tandem repeats, duplication regions, alignability smaller than 1 | 72 % | $\frac{nb_{de\ novo} \times SP\ \div SE}{2\ \times CG}$ SP : specificity SE : sensitivity |
| **Feng et al. 2017** | Herring | 12 | PCR free | Illumina HiSeq 2500 | 65.8X | BWA 0.6.2 | - | | GATK 3.3.0 - HaplotypeCaller and Samtools 1.19 mpilup | Customized filtering, overlapping of variant caller | Sanger sequencing | | Simulation and detection of true heterozygotes | 5.9 % | Remove site with mappability lower than 1 and repeat region | 51 % | $\frac{nb_{de\ novo\ all\ trios}}{2\ \times nb_{trios} \times CG \times (1-FDR)}$ |
| **Harland et al. 2017** | Cattle | 5 | PCR free | Illumina HiSeq 2000 | 23X | BWA mem 0.7.9a -r786 | Picard MarkDuplicates | Yes | GATK 3.4 - HaplotypeCaller | Customized filters | Illumina resequencing | 3% | | | Sites passing the DP filters | | |
| **Jónsson et al. 2017** | Human | 1550 | PCR free and non PCR free | Illumina | 35X | BWA mem 0.7.10 | Picard MarkDuplicates 1.117 | | GATK-UnifiedGenotyper | Customized filtering | Discordance monozygotic twin | 3% | Simultation of SNPs | 3.86 % | Sites with DP > 10 X and < 120 X (only reads MQ > 20, windows of 10,000 bp) | 88 % | $\frac{nb_{de\ novo}}{2\ \times CG}$ |

| | | | | | | | | | | | | | | | | | |
|---|---|---|---|---|---|---|---|---|---|---|---|---|---|---|---|---|---|
| **Maretty et al. 2017** | Human | 150 | | Illumina HiSeq 2000 | 78X | BWA mem 0.7.5a | yes | Yes | GATK - HaplotypeCaller | Customized filtering | | | | | 2.5 Gb | 87 % | |
| **Milholland et al. 2017** | Mouse | 8 | PCR free | Illumina HiSeq 2500 | 28.5X | BWA mem | Samtools | Yes | GATK - Unifiedgenotyper | DeNovoGear and VarScan2, kept overlapping candidates | Validation Sanger sequencing | 25 % | | | Number of base callable | 57.4 % | $\frac{nb_{de\ novo} \times (1-FDR)}{2 \times CG}$ |
| **Pfeifer 2017** | African green monkey | 3 | | Illumina HiSeq 2000 | 22X | BWA mem 0.7.13 | Picard MarkDuplicates 2.1.1 | Yes | GATK 3.5 - HaplotypeCaller | Customized filtering | Manual curation | 91 % | Mutations simulation | 15.6 % | Removed sites with $DP_{parent} < 10$, LowQual, MQ < 60, in repetitive regions, or incomplete genotyping | 57 % | $\frac{nb_{de\ novo}}{2 \times CG \times (1-FNR)}$ |
| **Tatsumoto et al. 2017** | Chimpanzee | 1 | PCR free | Illumina HiSeq 2000 | 150X | BWA mem 0.6.1 | Picard MarkDuplicates 1.93 | Yes | GATK 2.1.9 - UnifiedGenotyper | Customized filtering | Sanger validation | 20 % | DeNovoGear and validation | 0 | Remove sites with DP < filter | 45 % | $\frac{nb_{de\ novo} \times (1-FDR)}{2 \times CG}$ |
| **Turner et al. 2017** | Human | 516 | PCR free | Illumina HiSeq X | 34.8X | BWA mem 0.7.8 | Picard MarkDuplicates 1.83 | Yes | GATK 3.5 - HaplotypeCaller and FreeBayes | Customized filters and overlapping between varian | resequencing Sanger validation | 3.7 % | | | | | |

| | | | | | | | | | | | | | | | | | |
|---|---|---|---|---|---|---|---|---|---|---|---|---|---|---|---|---|---|
| | | | | | | | | | 1.0.1 | t caller | | | | | | | |
| **Malinski et al. 2018** | Cichlid (3 species) | 9 | | Illumina HiSeq 2000 | 40X | BWA mem 0.7.10 | Picard MarkDuplicates | | GATK 3.3.0 - HaplotypeCaller and samtools/bcftools 1.18 | Customized filters and overlapping between variant caller | From the binomial distribution of the AB | 5% | Read Position or Base Quality rank-sum test | 7.17 % | Using GATK filters on non variant site | 91.5 % | $\frac{nb_{de\ novo}}{2\ \times CG}$ |
| **Martin et al. 2018** | Platypus | 2 | | Illumina HiSeq 2000 | 16X | Stampy | Picard MarkDuplicates | | PLATYPUS | PLATYPUS *de novo* variant caller | | | | | | 46 % | $\frac{nb_{de\ novo\ 1} +\ nb_{de\ novo\ 2}}{CG_{trio\ 1} + CG_{trio\ 2}}$ |
| **Thomas et al. 2018** | Owl monkey | 14 | PCR free | Illumina HiSeq X | 37X | BWA mem 0.7.12 | Picard MarkDuplicates 1.105 | | GATK 3.3.0 - HaplotypeCaller | Customized filtering | Strict filters | 0 | correction for AB filter | 44 % | | | $\frac{nb_{de\ novo}}{2\ \times CG\ \times (1 - FNR)}$ |
| **Besenbacher et al. 2019** | Chimpanzee orangutan gorilla | 7,1,2 | PCR free | Illumina HiSeq X | 42X | BWA mem | | Yes | GATK 3.8 - HaplotypeCaller | Customized filtering | Strict filters | 0 | Correction for sites filters | | Probability of being a certain genotype given the depth | 32 - 79 % | $\frac{nb_{de\ novo}}{2\ \times CG\ \times (1 - FNR)}$ |
| **Koch et al. 2019** | Wolf | 4 | | Illumina HiSeq | 23X | BWA 0.7.12 | GATK 3.5.0 | Yes | GATK 3.5.0 - | Customized filteri | | | FNR as a probability | 11.2 % | Filters on a subset of sites | | Poisson distribution |

| | | | | | | | | | | | | | | | | | |
|---|---|---|---|---|---|---|---|---|---|---|---|---|---|---|---|---|---|
| | | | | 2000 | | | | | UnifiedGenotyper | ng | | | | | | | |
| **Lindsay et al. 2019** | Mouse | 15 | | Illumina HiSeq | 25X | | | | Bcftools and samtools | DeNovoGear 0.5 | Illumina resequencing | | | | Site passing DP filter and DeNovoGear prior filters | 89.9 % | |
| **Sasani et al. 2019** | Human | 593 | NonPCR free | Illumina HiSeq X | 30X | BWA mem 0.7.15 | Samblaster | Yes | GATK 3.5.0 | Customized filters | WGS resequenging higher depth and transmission | 4.5 % | "Missed heterozygote" also with transmition | 0.4 % | mosdepth to calculate DP and remove lower than 12, remove low MQ < 20, remove low complexity regions | 83.3 % | $\mu \times \frac{1 - FDR}{1 - MHR}$ |
| **Kessler et al. 2020** | Human | 1449 | PCR free | Illumina HiSeq X | 38X | BWA mem | | Yes | GotCloud pipeline | Customized filter and TrioDeNovo | call variant with TrioDeNovo | 1.8 % | Used similar FNR than Jonsson et al | 3.86 % | Site passing DP and quality threshold | | $\mu \times \frac{1 - FDR}{1 - MHR}$ |
| **Wang et al. 2020** | Rhesus macaque | 14 | PCR free | Illumina HiSeq X | 40X | BWA mem 0.7.12 | Picard Mark Duplicates | | GATK 3.6 | Customized filtering | | 0 | Probability | 20 % | Pass depth and homozygote reference filter | 88 % | $\frac{nb_{de\ novo}}{2 \times CG \times (1 - FNR)}$ |
| **Wu et al. 2020** | Baboon | 12 | PCR free | Illumina HiSeq X and Illumina HiSeq | 45X | BWA mem 0.7.9a | Samblaster 0.1.24 Picard Mark Duplicates | Yes | GATK 3.7 - HaplotypeCaller and | Customized filtering | Binomial probability of transmissi | 18 % | Mutations simulation | 7.93 % | GATK CallableLoci tool (on depth) + remove if Alt in parents | | Per sex likelihood |

| | | | | | | | | | | | | | | | | | |
|---|---|---|---|---|---|---|---|---|---|---|---|---|---|---|---|---|---|
| | | | | 2500 | | | 2.9.0 | | GenotypeGVCFs | | on | | | | | | |
| **Bergeron et al. 2021** | Rhesus macaque | 19 | NonPCR free | BGIseq500 | 81X | BWA mem 0.7.15 | Picard MarkDuplicates 2.7.1 | | GATK 4.0.7.0 - HaplotypeCaller BP_resolution | Customized filtering | Manual curation | 11 % | Correction for sites filters and AB filter | 4.28 % | Sites that pass HomRef, DP, and GQ filter | 89 % | $\frac{nb_{de\ novo} \times (1 - FDR)}{2\ \times CG\ \times (1 - FNR)}$ |
| **Campbell et al. 2021** | Grey mouse lemur | 2 | | 10X genomics | 34.5X | 10x Genomic's Long Ranger v2.2.1 | | | GATK 3.8 within Long Ranger | DeNovoGear v1.1.1 and VarScan2 v2.4.3, kept overlapping candidates | Error rate from technical replicates | 4.8 % | Error rate from technical replicates | 28 % | Simulation with BAMsurgeon v1.0.0 | 84 % | $\frac{nb_{de\ novo}}{2\ \times CG}$ |
| **Wang et al. 2021** | Cat | 11 | PCR free | Illumina HiSeq X | 41X | BWA mem 0.7.12 - r1039 | Picard MarkDuplicates 1.105 | | GATK 3.6 - HaplotypeCaller | Customized filters | | | On the AB filter | | Probability and correction for FNR | 72 % | $\frac{nb_{de\ novo}}{2\ \times CG\ \times (1 - FNR)}$ |
| **Yang et al. 2021** | Marmoset | 1 | NonPCR free | Offspring 10X genomics | Offspring 40X, Parents 75X | BWA ALN (v.0.7.12) | Picard MarkDuplicates | | GATK 4.0.7.0 - Haplo | Customized filtering | Validation with the other | | Correction for sites filters and AB filter | | Sites that pass HomRef, DP, GQ, and AD filter | 41 % | $\frac{nb_{de\ novo\ M}\ +\ nb_{de\ no}}{CG_M(1 - FNR_M) +\ CG_F(1}$ <br>M : mother<br>F : father |

| | | | | | | | | | | | |
|---|---|---|---|---|---|---|---|---|---|---|---|
| | | | | and parent Illumina | | | | | typeCaller BP_resolution | | haplotype (maternal and paternal) |

**Supplementary Table 2 – Description of site filters included in the GATK Best Practices hard-filtering recommendations.** See GATK https://gatk.broadinstitute.org/hc/en-us/articles/360035890471-Hard-filtering-germline-short-variants for more details

| **Site filter** | **Explanation** |
|---|---|
| **QualByDepth (QD)** | The variant confidence (QUAL field) divided by the unfiltered depth of non-HomRef samples. Used to select high-quality calls, and correct for the depth differences between samples. It is advised by GATK to use QD instead of the depth or the QUAL directly. |
| **RMSMappingQuality (MQ)** | Root mean square of the mapping quality over all the reads at the site. Good mapping qualities are around 60. Using the root square includes the standard deviation. High standard deviations mean that reads are highly divergent from the mean in this location. |
| **FisherStrand (FS)** | The Phred-scaled probability of strand bias at the site. Used to remove sites where the alternative allele is only seen in either forward or reverse strands. FS = 0 means no bias. |
| **StrandOddsRatio (SOR)** | FS penalizes variants at the end of exons, thus, SOR is another way to estimate and correct for strand bias. |
| **MappingQualityRankSumTest (MQRankSum)** | Compares the mapping qualities of the reads with the reference allele with those of the alternate allele and corrects for any bias. Positive values indicate a better mapping quality for alternative alleles, negative values indicate a bias toward the mapping quality of reference allele, and 0 means no bias. |

| | |
|---|---|
| **ReadPosRankSumTest (ReadPosRankSum)** | Compares the positions of the reference and alternate alleles on the reads. Negatives values indicate that alternative alleles are at the end of reads more often than reference alleles. Positive values indicate the opposite while 0 indicates no position bias. |

**Supplementary Table 3 – Several of the customized individual filters applied in the detection of germline mutations in the different reviewed studies.**

| Studies | Variant calling | ***Site filters (GATK recommended filters: QD* < 2.0 ; *MQ* < 40.0; *FS* > 60.0; *SOR* > 3.0; *MQRankSum*< -12.5; *ReadPosRankSum* < -8.0)** | Depth | Genotype quality | Allelic depth parents | Allelic balance |
|---|---|---|---|---|---|---|
| **Smeds et al. 2016** | GATK 3.3.0 - HaplotypeCaller and GenotypeGVCFs | | DPmin none DPmax none | GQ < 30 | AD > 0 | AB < 0.25 |
| **Harland et al. 2017** | GATK HaplotypeCaller (version 3.4) | | DPmin 10 DPmax 60 | GQ < 40 (offspring) | AD > 0 | |
| **Jónsson et al. 2017** | GATK-UnifiedGenotyper | "GATK best practice" | DPmin 12 DPmax none | GQ < 20 | AD > 1 | AB < 0.25 AB > 0.75 |

| | | | | | | |
|---|---|---|---|---|---|---|
| **Maretty et al. 2017** | GATK - HaplotypeCaller (gvcf) | “GATK best practice” | DPmin 20 DPmax 150 | $GQ_{Hom}$ < 80 $GQ_{Het}$ < 250 | AD > 4 | AB < 0.3 |
| **Pfeifer 2017** | GATK 3.5 - HaplotypeCaller and GenotypeGVCFs | LowQual, MQ < 60 | $DP_{parent}$ < 10 | None | AD > 0 | AB < 0.2 |
| **Tatsumoto et al. 2017** | GATK 2.1.9 - UnifiedGenotyper | | DPmin $av_i - 3\sigma$ DPmax $av_i + 3\sigma$ | $GQ_{Hom}$ < 100 $GQ_{Het}$ < 200 | None | At least 1 Read alt forward and 1 alt reverse |
| **Thomas et al. 2018** | GATK 3.3.0 - HaplotypeCaller | QD < 2.0; MQ < 40.0; FS > 60.0; ~~**SOR > 3.0**~~; MQRankSum < -12.5; ReadPosRankSum < -8.0 | DPmin 20 DPmax 60 | None | None | AB < 0.4 AB > 0.6 |
| **Besenbacher et al. 2019** | GATK 3.8 - HaplotypeCaller | FS > 20; ReadPosRankSum < -6 and > 6; BaseQualityRankSum; MappingQualityRankSum < 6 | DPmin 10 DPmax 1.9 × $av_i$ | GQ < 65 | AD > 0 lowQ AD2 > 1 | AB < 0.3 |
| **Koch et al. 2019** | GATK 3.5.0 - UnifiedGenotyper | QD < 2.0; MQ < 40.0; FS > 60.0; ~~**SOR > 3.0**~~; MQRankSum < -12.5; **ReadPosRankSum < 15** | DPmin 10 DPmax 100 | None | AD > 0 | At least 1 read with Alt |
| **Sasani et al. 2019** | GATK v3.5.0 | | DPmin 12 DPmax none | GQ < 20 | AD > 0 | More than 0.3% of the reads for the 3rd generation (transmission was not possible) |
| **Wang et al. 2020** | GATK 3.6 | QD < 2.0; MQ < 40.0; FS > 60.0; ~~**SOR > 3.0**~~; MQRankSum < -12.5; ReadPosRankSum < -8.0 | DPmin 20 DPmax 60 | GQ < 70 | AD > 2 | AB < 0.35 |

| | | | | | | |
|---|---|---|---|---|---|---|
| **Wu et al. 2020** | GATK 3.7 - HaplotypeCaller and GenotypeGVCFs | QD < 2.0 ; MQ < 40.0; FS > 60.0; SOR > 3.0; MQRankSum< -12.5; ReadPosRankSum < -8.0 | pvalue poisson test $2 \times 10^{-4}$ | GQ < 40 | AD = 0 at least for one parent | At least 3 reads with Alt and binomial test on allelic balance p-value < 0.05 |
| **Bergeron et al. 2021** | GATK 4.0.7.0 - HaplotypeCaller | QD < 2.0; MQ < 40.0; **FS > 20.0**; SOR > 3.0; **MQRankSum < -2.0; MQRankSum > 4.0; ReadPosRankSum < -3.0; ReadPosRankSum > 3.0** | DPmin $0.5 \times av_t$ DPmax $2 \times av_t$ | GQ < 60 | None | AB <0.3 AB > 0.7 |
| **Wang et al. 2021** | GATK version 3.6 | QD < 2.0; MQ < 40.0; FS > 60.0; ~~**SOR > 3.0**~~; MQRankSum < -12.5; ReadPosRankSum < -8.0 | DPmin 20 DPmax 60 | GQ < 70 | AD > 0 | AB < 0.35 |

**Supplementary Table 4 – Methodology and filtering criteria of the five different pipelines applied on the common dataset**.

| Step of the analysis | Information to report | LB pipeline | TT pipeline | RW pipeline | CV pipeline | SB pipeline |
|---|---|---|---|---|---|---|
| **1 – Sampling and sequencing** | Type of sample | Whole blood stored in EDTA | Same as LB for Mutationathon | Same as LB for Mutationathon | Same as LB for Mutationathon | Same as LB for Mutationathon |
| | Age of sample | Freshly collected, stored at -80°C, and ship in dry ice. | Same as LB for Mutationathon | Same as LB for Mutationathon | Same as LB for Mutationathon | Same as LB for Mutationathon |
| | Type of library preparation | Libraries were not PCR-free | Same as LB for Mutationathon | Same as LB for Mutationathon | Same as LB for Mutationathon | Same as LB for Mutationathon |
| | Average coverage | Father 40 X and mother and offspring 70 X | Same as LB for Mutationathon | Same as LB for Mutationathon | Same as LB for Mutationathon | Same as LB for Mutationathon |
| | Autosomes only or with sex chromosomes ? | Autosome only | Autosomes and sex chomosomes | Autosome only | Autosome and sex chromosomes | Autosome only |
| **2 – Alignment and post-alignment processing** | Trimming of adaptors and low quality reads | Yes with SOAPnuke | Same as LB for Mutationathon | Same as LB for Mutationathon | Same as LB for Mutationathon | Same as LB for Mutationathon |
| | Which reference assembly | Mmul 8.0.1 | Mmul 8.0.1 | Mmul 8.0.1 | Mmul 8.0.1 | Mmul 8.0.1 |
| | Mapping software and version | BWA mem 0.7.15 with insert size option | BWA mem 0.7.17-r1188 | BWA-MEM v. 0.7.12 | BWA mem 0.7.15 | BWA mem 0.7.15 |

| | | | | | | |
|---|---|---|---|---|---|---|
| | Removing duplicates software and version | Removed with Picard 2.7.1 | Marked with Picard 2.21.4 | Picard MarkDuplicates v. 1.105 | Picard MarkDuplicates v. 2.18.3 | Samtools 1.6.0 |
| | Base quality score recalibration (yes/no) | No | No | No | Yes | Yes |
| | If yes, which type of data used as known variants | N/A | N/A | N/A | dbSNP build 150 | Harris et. al, 2020 (https://doi.org/10.1186/s12862-020-1595-9) |
| | Realignments around indels? | No | No | No | No | Yes |
| | Other filters? | Removing reads mapping to multiple locations of the genome with samtools 0.1.18 | No | No | 2nd round of Picard MarkDuplicates v. 2.18.3 (post-BQSR) | Use GATK PrintReads with the following filters: BadCigar, DuplicateRead, FailVendorQualityCheck, HCMappingQuality, MappingQualityUnavailable, NotPrimaryAlignment, UnmappedRead, filter_bases_not_stored, filter_mismatching_base_and_quals |
| **3 – Variant calling** | Software and version | GATK 4.0.7.0 | GATK 3.5.0, freebayes v1.3.2-40-gcce27fc, Platypus2 v 0.8.1.1, | GATK 4.1.7.0 | GATK 3.7 | GATK 3.8 |

<table>
<tr><td></td><td></td><td></td><td>Strelka2 v 2.9.10</td><td></td><td></td><td></td></tr>
<tr><td></td><td>Mode: joint genotyping? Gvcf blocks? Gvcf in base-pair resolution?</td><td>HaplotypeCaller in BP-RESOLUTION per individuals; combined individuals with CombineGVCFs; joint genotyping with GenotypeGVCF</td><td>joint-genotyping across crams</td><td>HaplotypeCaller, joint call with GenotypeGVCF, followed by CalculateGenotypePosterior.<br>Not bp resolution.</td><td>HaplotypeCaller in BP-RESOLUTION; joint genotyping with GenotypeGVCF</td><td>HaplotypeCaller with all sample at once</td></tr>
<tr><td>4 – Detecting de novo mutations</td><td>Site filters on vcf files and justification</td><td>QD < 2.0, FS > 20.0, MQ < 40.0, MQRankSum < - 2.0, MQRankSum > 4.0, ReadPosRankSum < -3.0, ReadPosRankSum > 3.0 and SOR > 3.0. The filters were chosen to reduce false-positive calls after a first manual curation of the candidates using the recommended filters.</td><td>Mother genotype = 0/0, Father genotype = 0/0, Child genotype = 0/1 or 1/1 or 1/0 or 0|1 or 1|1 or 1|0. Remove variants in recent repeats. Remove variants in homopolymers of AAAAAAAAAA or TTTTTTTTTT</td><td>QD < 2.0 || MQ < 40.0 || FS > 60.0 || SOR > 3.0 || MQRankSum < -12.5 || ReadPosRankSum < -8.0<br><br>Our standard set of hard filters for genotype calling.</td><td>autosomal biallelic SNPs fully genotyped in the extended trio, GATK Best Practices hard filter criteria</td><td>FS > 30.0, MQRankSum < - 10, MQRankSum > 10, ReadPosRankSum < -2.5, ReadPosRankSum > 2.5, BaseQRankSum < -13, BaseQRankSum >13</td></tr>
</table>

| | | | | | | |
|---|---|---|---|---|---|---|
| | Individual filters, threshold, and remaining candidates after each filter | Mendelian violation parents HomRef and offspring Het (11,779 candidates); AB: 0.3 and 0.7 (candidates 5,527); DP : 0.5 × depth individual and 2 × depth individual ; 20 X to 78X for the father, 30X to 121X for the mother, and 31X to 125X for the offspring ( 951 candidates); GQ: 60 (34 candidates); AD: none | No evidence of alternate allele in Father, No evidence of alternate allele in mother, AB in child > 0.25, depth in each individual >= 10, genotype quality > 20 | Mendelian violations where parents were HomRef and offspring was Het.<br><br>20 < DP < 80<br>Alt AB > 0.35<br>GQ > 20<br>Alt AD < 1 in parent<br>Alt ADF > 0, ADR > 0 in offspring | Mendelian violation where parents were HomRef and offspring was Het<br><br>0.5 × $DP_{ind}$ & 2 × $DP_{ind}$; 0.25<=AB<=0.75; GQ>=40;<br>$AD_{alt}$ = 0 in parents | Mendelian violation, parents HomRef and offspring Het; AB: >0.3; DP: min 10 and max 1.75x median of individual; GQ: min 55; no good qual(>20) alt reads in parents and max 1 bad qual(<20) read; alt allele should be seen on both strands in child |
| | False discovery rate estimation method: PCR validation? Manual curation? Transmission rate deviation? | Manual curation with IGV and removed 6 candidates (28 final candidates) | Since there was no low-complexity region (LCR) annotation file for rheMac8, I did do a samtools tview assessment of all candidate DNVs. After removing all DNVs in LCRs there were 34 high-confidence DNVs (29 SNVs [2 at CpG sites], 4 INDELS, 1 MNV) | Manual curation with IGV, but no candidates were removed.<br>High transmission rate noted (72%), but not used to adjust rate estimate. | - | Assumes filters are set so conservatively that no false positives remain |

| | | | | | | |
|---|---|---|---|---|---|---|
| **5 – Mutation rate estimation** | Method to access callability: File used? Filters took into account? | All sites passing the DP and GQ filters on the BP resolution vcf files (including all sites in the genome). | For mutation rate estimation, I assessed the length of the genome without gaps or other regions excluded from variant calling (ungapped length = 3142076610 bp). Only assessed SNVs for the mutation rate estimation. For me it was 29 SNVs / (3142076610 bp *2) = 0.46 x 10-8 substitutions per site per generation | Our equivalent here would be the enumeration of sites from samtools mpileup that pass the DP filter | applied the same filters as for DNM detection on the BP resolution .vcf (including all sites in the genome) | The callability of each site is inferred as a probability conditional on the depth in each of member of the trio. These probabilities are then summed. See Besenbacher 2015, Nat. Comm. |
| | False-negative rate estimation method: simulation? Filters? Probability? | Proportion of sites expected to be filtered away by the 2 remaining filters: the site filters (following known distributions) and the allelic balance filter (proportion of true heterozygotes outside AB filters). | - | Our equivalent estimate of FNR would be the product of parental homozygote callability and offspring heterozygous callability. We sample 250k high-quality SNPs for each of the two categories and subject them respectively to the parental and offspring filters used for the de novo candidates. The proportion of remaining SNPs after all filtering | - | The false negative rate of all filters except the site filters are incorporated in the number callable sites estimates as explained above. The sites filters used follow a known null distribution an we use correct for the expected false negative rates of those. |

| | | | | |
|---|---|---|---|---|
| | | | | are interpreted as the probability that a true de novo candidate would pass the filter. |

**Supplementary Table 5 – PCR validation of DNM candidates found by the various pipelines during the Mutationathon.** TP means validated as true positive DNM and FP appeared as false positive. The genotypes of all individuals as shown by the PCR validation are presented.

| **Chromosome** | **Position** | **Ref** | **Alt** | **PCR validation** | **Genotypes** | | | | **Transmission** | **Pipelines** | | | | | |
|---|---|---|---|---|---|---|---|---|---|---|---|---|---|---|---|
| | | | | | **Heineken (offspring)** | **Noot (father)** | **M (mother)** | **Hoegaarde (2nd offspring)** | | **CV** | **RW** | **TT** | **LB** | **SB** | **Number of pipelines** |
| **chr2** | 5858286 | A | G | FP | A/A | A/A | A/A | A/A | | x | | | | | 1 |
| **chr2** | 60917810 | A | G | not amplified | | | | | | | | x | | | 1 |
| **chr2** | 150987129 | A | G | TP | A/G | A/A | A/A | A/A | no | x | | x | x | x | 4 |
| **chr2** | 203383064 | C | T | TP | C/T | C/C | C/C | C/C | no | x | | x | x | x | 4 |
| **chr3** | 76549728 | G | T | TP | G/T | G/G | G/G | T/T | no | | x | | x | x | 3 |
| **chr3** | 94315937 | C | G | TP | C/G | C/C | C/C | C/G | yes | | x | | | x | 2 |
| **chr4** | 116799040 | G | A | TP | G/A | G/G | G/G | G/A | yes | x | x | x | x | x | 5 |
| **chr5** | 85980247 | A | G | FP | A/A | A/A | A/A | A/A | | x | | | | | 1 |
| **chr5** | 118992462 | T | C | TP | C/T | T/T | T/T | T/C | yes | | x | x | x | x | 4 |
| **chr6** | 11241018 | A | G | not amplified | G/A | | A/A | A/G | | | x | x | x | x | 4 |
| **chr6** | 18650255 | G | A | TP | A/G | G/G | G/G | A/G | yes | x | x | | x | | 3 |
| **chr6** | 64657185 | C | T | TP | C/T | C/C | C/C | C/T | yes | | | | x | | 1 |
| **chr6** | 157547189 | G | A | TP | A/G | G/G | G/G | G/G | no | x | | x | x | x | 4 |
| **chr7** | 149216879 | G | A | TP | G/A | G/G | G/G | G/A | yes | x | x | x | x | x | 5 |
| **chr7** | 153210388 | T | C | TP | T/C | T/T | T/T | T/C | yes | x | x | x | x | x | 5 |
| **chr8** | 7226159 | G | T | FP | G/G | G/G | G/G | G/G | | | | | | x | 1 |
| **chr8** | 7226160 | G | T | FP | G/G | G/G | G/G | G/G | | | | | | x | 1 |
| **chr8** | 11458034 | C | T | TP | C/T | C/C | C/C | C/C | no | | | x | x | x | 3 |

| | | | | | | | | | | | | | | | |
|---|---|---|---|---|---|---|---|---|---|---|---|---|---|---|---|
| **chr9** | 332616 | G | A | TP | A/G | G/G | G/G | A/G | yes | | x | x | x | x | 4 |
| **chr9** | 35992733 | G | T | TP | T/G | G/G | G/G | T/G | yes | x | | x | x | x | 4 |
| **chr9** | 36684503 | T | A | TP | T/A | T/T | T/T | T/A | yes | x | x | x | | x | 4 |
| **chr9** | 57966486 | T | C | TP | T/C | T/T | T/T | T/T | no | | x | x | x | x | 4 |
| **chr9** | 58787884 | T | C | not amplified | | | | | | | | | x | | 1 |
| **chr9** | 59187032 | T | C | TP | T/C | T/T | T/T | T/T | no | x | x | x | x | x | 5 |
| **chr9** | 88707756 | G | A | TP | A/G | G/G | G/G | A/G | yes | x | x | x | x | x | 5 |
| **chr9** | 93536046 | C | T | TP | T/C | C/C | C/C | C/C | no | | | | x | | 1 |
| **chr9** | 125284007 | G | A | TP | A/G | G/G | G/G | G/G | no | | x | x | x | x | 4 |
| **chr10** | 40351695 | C | A | TP | C/A | C/C | C/C | C/A | yes | x | x | x | x | x | 5 |
| **chr10** | 65456149 | G | A | TP | G/A | G/G | G/G | G/G | no | | x | x | | x | 3 |
| **chr11** | 44403758 | G | A | TP | A/G | G/G | G/G | A/G | yes | | | | x | | 1 |
| **chr11** | 79918225 | A | C | TP | A/C | A/A | A/A | A/C | yes | | | | | x | 1 |
| **chr11** | 98692026 | G | A | TP | G/A | G/G | G/G | G/G | no | | x | x | x | x | 4 |
| **chr11** | 109301631 | C | T | TP | C/T | C/C | C/C | C/C | no | x | x | x | x | x | 5 |
| **chr12** | 110983332 | C | G | TP | C/G | C/C | C/C | C/G | yes | | | x | x | x | 3 |
| **chr13** | 28463207 | C | T | TP | T/C | C/C | C/C | T/C | yes | | x | x | x | x | 4 |
| **chr13** | 79024526 | C | A | TP | A/G | C/G | G/G | A/G | yes | | | x | | | 1 |
| **chr13** | 105761009 | T | G | FP | G/G | G/G | G/G | G/G | | | | | | x | 1 |
| **chr14** | 41517101 | T | C | TP | T/C | T/T | T/T | T/C | yes | x | x | x | x | | 4 |
| **chr14** | 52516157 | C | G | TP | G/C | C/C | C/C | G/C | yes | x | x | x | | x | 4 |
| **chr15** | 27373968 | C | T | TP | T/C | C/C | C/C | T/C | yes | x | x | | x | x | 4 |
| **chr16** | 22611461 | C | T | TP | T/C | C/C | C/C | T/C | yes | | | x | | | 1 |
| **chr18** | 38858125 | T | C | FP | C/T | C/T | T/T | T/T | | | | | | x | 1 |
| **chr19** | 16726793 | G | A | not amplified | | | | | | | | | | x | 1 |

**Supplementary Table 6 – The impact of individual filters on the estimated rate of the trio of rhesus macaques**.

| **Individual filters** | **Candidate DNMs** | **False-positive calls** | **Callable genome** | **FNR** | **Mutation rate per site per generation** |
|---|---|---|---|---|---|
| DP < 10 | 46 | 16 | 2,326,488,078 | 0.042 | $0.67 \times 10^{-8}$ |
| DP < 12 | 46 | 16 | 2,326,488,069 | 0.042 | $0.67 \times 10^{-8}$ |
| DP < 20; DP > 150 | 43 | 14 | 2,325,499,068 | 0.041 | $0.65 \times 10^{-8}$ |
| DP < 20; DP > 80 | 43 | 13 | 1,788,223,190 | 0.423 | $0.67 \times 10^{-8}$ |
| DP < 0.5 × $dp_{ind}$; DP > 2 × $dp_{ind}$ | 34 | 6 | 2,312,067,783 | 0.040 | $0.63 \times 10^{-8}$ |
| No GQ | 951 | 870 | 2,371,357,494 | 0.041 | $1.8 \times 10^{-8}$ |
| GQ < 20 | 65 | 24 | 2,355,822,929 | 0.040 | $0.91 \times 10^{-8}$ |
| GQ < 40 | 42 | 11 | 2,346,070,553 | 0.040 | $0.69 \times 10^{-8}$ |
| GQ < 60 | 34 | 6 | 2,312,067,783 | 0.040 | $0.63 \times 10^{-8}$ |
| GQ < 80 | 23 | 0 | 2,147,415,676 | 0.039 | $0.56 \times 10^{-8}$ |
| $GQ_{Hom}$ < 100 and $GQ_{Het}$ < 200 | 20 | 0 | 1,712,742,807 | 0.038 | $0.61 \times 10^{-8}$ |
| No AD | 34 | 6 | 2,312,067,783 | 0.040 | $0.63 \times 10^{-8}$ |
| AD > 0 in at least 1 parent | 34 | 6 | 2,303,628,249 | 0.040 | $0.63 \times 10^{-8}$ |
| AD > 4 | 33 | 5 | 2,312,079,962 | 0.040 | $0.63 \times 10^{-8}$ |
| AD > 1 | 32 | 4 | 2,300,625,789 | 0.040 | $0.63\times 10^{-8}$ |
| AD > 0 | 28 | 3 | 2,088,477,209 | 0.040 | $0.62 \times 10^{-8}$ |
| AB < 0.2 | 52 | 15 | 2,312,067,783 | 0.035 | $0.83 \times 10^{-8}$ |
| AB < 0.25 | 38 | 9 | 2,312,067,783 | 0.036 | $0.65 \times 10^{-8}$ |
| AB < 0.25; AB > 0.75 | 38 | 9 | 2,312,067,783 | 0.036 | $0.65 \times 10^{-8}$ |
| AB < 0.3 | 34 | 6 | 2,312,067,783 | 0.040 | $0.63 \times 10^{-8}$ |
| AB < 0.3; AB > 0.7 | 34 | 6 | 2,312,067,783 | 0.040 | $0.63 \times 10^{-8}$ |
| AB < 0.35 | 31 | 5 | 2,312,067,783 | 0.061 | $0.60 \times 10^{-8}$ |
| AB < 0.4; AB > 0.6 | 27 | 3 | 2,312,067,783 | 0.158 | $0.62 \times 10^{-8}$ |

**Supplementary Table 7 – PCR experiment and Sanger resequencing.** Primers for the 40 positions validated with PCR experiment. For each individual, the GenBank accession number are indicated as sequenceID with first the forward strand and then the reverse.

| Chromosome | Position | Forward Primer sequence (5′-3′) | Reverse Primer sequence (5′-3′) | SequenceID father F/R | SequenceID mother F/R | SequenceID offspring F/R | SequenceID 2nd offspring F/R |
|---|---|---|---|---|---|---|---|
| chr2 | 5858286 | CTGAATTAGCCCACACTACA | CATGGAGTCACATTAAGGCT | EF31864958/- | -/EF31864936 | EF31877336/- | EF31864919/EF31864921 |
| chr2 | 150987129 | CATTAAGCCCCAGGTACATT | TTTTCCATAAATGGGCACCT | EF31873919/- | EF31872229/EF31872230 | EF31864006/EF31864007 | EF31872228/EF31872315 |
| chr2 | 203383064 | GGGGCATCTTCATTTTTCAC | ATACCCTATGCCACTTTACC | EF31873922/EF31873923 | -/EF31872242 | EF31877338/EF31877339 | EF31872239/EF31872240 |
| chr3 | 76549728 | TCTTAAGCCTCCACCAATTT | TTGTCAACATTGGTTCAACA | EF31873924/EF31873925 | EF31872253/EF31872254 | EF31864010/EF31864011 | -/EF31872252 |
| chr3 | 94315937 | TCACCCCACTTGTATTATGG | | EF31864960/- | EF31864937/- | EF31864981/- | EF31864922/- |
| chr4 | 116799040 | GTTGTGGTCTGGGAGTATAA | GCTCCTACCTATTTGCTCTG | EF31873928/EF31873929 | EF31872277/EF31872278 | EF31864015/EF31864017 | -/EF31872276 |
| chr5 | 85980247 | GGAGACGCTTCTTATAACTCT | TACTGCTGCTCTCACTATC | EF31864961/EF31864962 | EF31864938/EF31864939 | EF31864593/EF31864594 | EF31864924/EF31864925 |
| chr5 | 118992462 | AAGCACTTGTAAGAGTACCCA | TCTCCCTGTAACATGACTTAAA | EF31873932/EF31873933 | EF31872303/EF31872304 | EF31864027/EF31864028 | EF31872300/EF31872302 |
| chr6 | 11241018 | CACAGGTAATAAAGAAGGGA | TTCCCACCAACAGTATAAAA | -/- | EF31872219/EF31872213 | EF31864093/EF31864094 | EF31873848/EF31873857 |
| chr6 | 18650255 | ATGCCAAGACTTGAGATGAG | GTCGTGACTGTGTTCTTAGT | EF31864581/EF31864582 | EF31872237/EF31872238 | EF31877378/EF31877379 | EF31873904/EF31873914 |
| chr6 | 64657185 | TCAAATTCATAGAGGCAGAA | ACAACCTGCTAAAAATGAAG | EF31864583/EF31864584 | EF31872249/EF31872250 | EF31877346/EF31877347 | -/EF31873915 |
| chr6 | 157547189 | CTTGTGTGGTATGCTTTATG | GATACCACAGATCTACAGAC | EF31873936/ | EF31872231/ | EF31864097/ | EF31873849/ |

| | | | | | | | |
|---|---|---|---|---|---|---|---|
| | | | | EF31873937 | EF31872232 | EF31864099 | EF31873858 |
| chr7 | 149216879 | CACAGTTGCTCTTCATTAGTC | CAGTCTGGAAACTCTCATCAT | EF31873938/ EF31873939 | EF31872243/EF31 872244 | EF31877382/ EF31877383 | EF31873850/EF31 873859 |
| chr7 | 153210388 | CCACCAAAATTGCTCTTGAT | CAGCTTGTTCAGGGTTACTC | EF31873940/ EF31873941 | EF31872255/EF31 872256 | EF31864033/ EF31864034 | EF31873851/EF31 873860 |
| chr8 | 7226159 and 7226160 | CGGTCATGTTAGGATGGAG | AAAGTACTTGAGTCCATGACA | EF31873942/ EF31873943 | EF31872267/EF31 872268 | EF31877384/ EF31877385 | EF31873852/EF31 873861 |
| chr8 | 11458034 | GCTTTAATTAGCTTCAGTCCG | CATTAATTTTGACTGCCATGTG | EF31873944/ EF31873945 | EF31872279/EF31 872281 | EF31877386/ EF31877388 | EF31873853/EF31 873863 |
| chr9 | 332616 | CACTCGATTAGCACCATTCT | GTGTTTCGAGCAATCTACAC | EF31873946/ EF31873947 | EF31872292/- | EF31864040/ EF31864041 | EF31873854/EF31 873864 |
| chr9 | 35992733 | | TAAAGTCTTGCCCTCAGTTC | - /EF31864586 | -/EF31872262 | - /EF31864043 | -/EF31873916 |
| chr9 | 36684503 | GAGCCATGTTCACGATTTTT (offspring:GATGTATGCTCTGAA CAGGA) | AATGGCTGAATAGCATACCC | EF31864587/ EF31864605 | EF31872273/EF31 872274 | EF31877391/ EF31864046 | EF31873908/- |
| chr9 | 57966486 | CTCACTAAAATGAGGGGACA | TAGTTTTTCTCACACCCACAT | EF31873948/ EF31873949 | EF31872305/EF31 872306 | EF31864047/ EF31864048 | EF31873855/EF31 873865 |
| chr9 | 59187032 | AGAAAGCAAGTGGTATGTGT | GTTGTAGTGTACAATCGCAT | EF31873950/ EF31873952 | EF31872214/EF31 872215 | EF31864051/ EF31864052 | EF31873866/EF31 873876 |
| chr9 | 88707756 | TCTATATGACCCCAGGATAG | AGGATGTCTGTTATTACCAC | EF31873953/ EF31873954 | EF31872233/EF31 872234 | EF31877393/ EF31877394 | EF31873867/EF31 873877 |
| chr9 | 93536046 | GGTCCCATTGGGGTAATATG | ATTGGTTGGCCTGAAGTTTA | EF31864546/ EF31864547 | EF31872245/EF31 872246 | EF31877350/ EF31877351 | EF31873868/EF31 873878 |
| chr9 | 125284007 | GAGAGGGCTAATTTGGAAGT | ACAAGGAGGTATTTCGGTTG | EF31864548/ EF31864603 | EF31872257/EF31 872258 | EF31877352/ EF31877353 | EF31873869/EF31 873879 |
| chr10 | 40351695 | ACATCTCCTTGACAGGATTG | TTGATGACCTGAGAAGTACC | EF31864550/ EF31864551 | EF31872269/EF31 872270 | EF31877354/ EF31877355 | EF31873870/EF31 873880 |
| chr10 | 65456149 | GTTGGGTATGAATGTGGACT | GCTCAGAACGTTTTTATGGA | EF31864552/ | EF31872282/EF31 | EF31864066/ | -/EF31873881 |

| | | | | | | | |
|---|---|---|---|---|---|---|---|
| | | | | EF31864554 | 872283 | EF31864067 | |
| chr11 | 44403758 | TACTGGGAACAGTACTGACA | ACTGGCTTGCTAAGGTTAAA | EF31864979/ EF31864980 | EF31872294/EF31 872295 | EF31877356/ EF31877357 | EF31873872/EF31 873882 |
| chr11 | 79918225 | | CCCAGCAATCTCTCTTCTTT | - /EF31873770 | -/EF31873769 | - /EF31873744 | -/EF31873768 |
| chr11 | 98692026 | AGAGCTATAGGAAGGCGATT | | EF31864589/ - | EF31872286/- | EF31864070/ - | EF31873909/- |
| chr11 | 109301631 | TTTTAAAGACAAGGTCTCGC | TATTTGCCAAACGTCTATTG | EF31864557/ EF31864559 | EF31872307/EF31 872308 | EF31864103/ EF31864104 | EF31873875/EF31 873883 |
| chr12 | 110983332 | TTTGGGGAGTGAGAGTAGAT | AACTGTCACATTCTCTCAGG | EF31864560/ EF31864563 | EF31872216/EF31 872224 | EF31864072/ EF31864073 | EF31873884/EF31 873893 |
| chr13 | 28463207 | GTCAACACCTTAGAAAAGGC | TGTACTCTGTAAGGAGACGA | EF31864564/ EF31864565 | EF31872235/EF31 872236 | EF31864074/ EF31864075 | EF31873885/EF31 873895 |
| chr13 | 79024526 | TATTGTAACACACAGGCCAA | GCTAAACGACATTTACCAGC | EF31864566/ - | EF31872247/EF31 872248 | EF31877369/ EF31877370 | EF31873886/- |
| chr13 | 105761009 | GCTAAAAACTCAATAAATTCG G | ATAAATTACCTTGGGCACTA | EF31864568/ EF31864569 | EF31872259/EF31 872260 | EF31864105/ EF31864106 | EF31873887/EF31 873898 |
| chr14 | 41517101 | GTCTTAGGCAGTGATTGTGA | GAGCAAGAGCATCATTGTTT | EF31864570/ - | EF31872271/EF31 872272 | EF31877401/ EF31877402 | EF31873888/EF31 873899 |
| chr14 | 52516157 | TTTGTGAGAACCCATCTCCG | | EF31873773/ - | EF31873772/- | EF31873745/ - | EF31873771/- |
| chr15 | 27373968 | GAAACACTGGAAGCACATAC | TTTGCACTCACTGATCAACT | EF31864573/ EF31864574 | EF31872284/EF31 872285 | EF31864078/ EF31864080 | EF31873890/EF31 873900 |
| chr16 | 22611461 | AAGTCTTAATACATTGGGGG | GCAATATGGCAAGAGATTG | - /EF31864965 | -/EF31864944 | EF31877407/ EF31864988 | EF31864927/- |
| chr18 | 38858125 | GAATGATTCAAGGCTGTTCTC | ATGGCCACTAAAAATCACTT | EF31864604/ EF31864578 | EF31872309/EF31 872310 | EF31877364/ EF31877365 | EF31873892/EF31 873902 |
